# Harnessing *Vitis* diversity to dissect and predict adventitious rooting traits in grapevine

**DOI:** 10.64898/2026.08.24.746882

**Authors:** Sadikshya Sharma, Yaniv Lupo, Jose Munoz, Noe Cochetel, Veronica Nunez, Ana Gaspar, Efrain Torres-Lomas, Dario Cantu, Luis Diaz-Garcia

## Abstract

Adventitious root formation is critical for the cost-effective propagation of grapevine rootstocks, and poor rooting limits the adoption of new rootstocks derived from underutilized *Vitis* L. species. We evaluated 308 accessions representing 18 *Vitis* species over three growing seasons, scoring rooting at two developmental stages, callus-stage and post-transplant, together with root biomass, cutting weight, and a derived transplant- response index. Phenotypic variation was extensive within and among species, and species rankings depended on the trait considered: *V. riparia*, *V. rupestris* and *V. californica* ranked among the top five species for all four rooting traits, whereas *V. arizonica* and *V. acerifolia* rooted well at the callus stage but only intermediately after transplanting. Repeatability was moderate to high for root weight and callus-stage rooting and lower for post-transplant rooting. Between-species differences accounted for most of the genetic variance in callus-stage rooting but little of that in cutting weight. After removing differences among species, accessions originating from wild sites with lower dry-season precipitation rooted better and produced more root biomass. Genome-wide association analysis of 3.4 million single-nucleotide polymorphisms identified 54 significant markers resolving into 18 independent loci across four traits. Candidate genes implicate auxin-linked cell proliferation, cell wall and lignin remodeling, and solute transport. Genomic and phenomic prediction achieved moderate accuracies across traits and seasons, including for previously unevaluated accessions, and combining spectral with genotypic data improved performance; accuracy was essentially flat between 5,000 and 50,000 markers. These results provide a framework for broadening the germplasm base of grapevine rootstock breeding.

**Plain Language Summary:** Grapevines are almost always grown as two plants joined together: a fruiting variety grafted onto a rootstock that supplies the root system. Nurseries build these plants from dormant cuttings, so a rootstock is only useful in practice if its cuttings root easily. Almost all commercial rootstocks descend from just three wild North American grape species, in part because cuttings of other species are believed to root poorly. We grew cuttings from 308 wild and cultivated grapevines representing 18 species over three years and measured how well each one rooted, first in the callusing room and again after the young plants were transplanted. Rooting ability varied a great deal, and several species outside the usual three rooted as well as the standards. Vines originally collected from places with drier summers tended to root best. We also located regions of the grape genome linked to rooting, and showed that the rooting ability of a vine can be predicted from its DNA or from light reflected by its leaves. Together, these results give breeders a way to screen a much wider range of wild grapevines for rootstock development before testing them in a nursery.

**Core Ideas:** Rooting ability varies widely across 18 *Vitis* species, well beyond the three used in rootstock breeding.

Callus-stage and post-transplant rooting behave as genetically distinct stages of root formation.

Accessions from collection sites with drier summers rooted better and produced more root biomass.

GWAS resolved 18 loci implicating auxin signaling, cell wall remodeling, and solute transport.

Genomic and phenomic prediction reached moderate accuracy and was insensitive to marker density.

## Introduction

The genus *Vitis* comprises approximately 70 interfertile species, with North America recognized as a center of origin due to its substantial species diversity (Wan et al. 2013). This genetic diversity has been instrumental to the development of rootstocks that sustain modern grapevine cultivation worldwide. Among these, *V. riparia*, *V. rupestris*, and *V. berlandieri*, and their interspecific hybrids, have played a central role. *Vitis riparia* and *V. rupestris* are valued for their robust rooting capabilities and phylloxera resistance (Pongrácz 1983; Viala & Ravaz 1903; Riaz et al. 2019), while *V. berlandieri* is primarily utilized for its tolerance to lime-induced chlorosis and adaptability to calcareous soils (Foëx 1902; Viala & Ravaz 1903). Beyond this trio, ARF in other *Vitis* species remains poorly characterized, limiting their use in rootstock breeding. Historically, rootstock selection has prioritized phylloxera resistance and lime tolerance (Pongrácz 1983; Viala & Ravaz 1903); however, current viticulture faces a broader spectrum of biotic and abiotic challenges. Soil-related stresses, including salinity, drought, heat, waterlogging, and poor drainage, can significantly impair vine performance, requiring rootstocks with specific adaptive traits (Lanyon et al. 2004; Oliver et al. 2013). Additionally, biotic threats beyond phylloxera, particularly plant-parasitic nematodes, cause substantial root damage, leading to vine decline and acting as vectors for viral infections (Ferris et al. 2012; McKenry & Anwar 2006). While the riparia-rupestris-berlandieri group has addressed many of these issues, the growing demand for sustainable agriculture, climate change pressures, and the emergence of new pests and diseases underscore the need to expand the genetic base of commercial rootstocks and enhance stress tolerance and resistance mechanisms.

Several under-utilized wild *Vitis* species offer promising potential for rootstock development, particularly those adapted to extreme environmental conditions. In the southwestern United States, species such as *V. arizonica*, *V. girdiana*, *V.* × *doaniana*, and *V. acerifolia* have evolved under harsh climates, developing unique adaptations beneficial for breeding. For example, certain accessions of *V. acerifolia* and *V.* × *doaniana* demonstrate exceptional tolerance to soil salinity, exhibiting high chloride exclusion capacity (Heinitz et al. 2020; Sharma et al. 2024; Sharma et al. 2025). Additionally, specific *V. arizonica* populations from the U.S. Southwest and northern Mexico possess strong genetic resistance to root pests and diseases, including root-knot nematodes (*Meloidogyne* spp.) (Askary et al. 2018), dagger nematodes (*Xiphinema index*) (Riaz et al. 2007), phylloxera (Johnson & Hilsenbeck 1989), and Pierce’s disease (Riaz et al. 2007). Assessing the ARF capacity of these and other wild *Vitis* species is a critical step toward integrating them into rootstock breeding programs and developing molecular breeding strategies to accelerate breeding cycles.

Despite extensive research on the physiological and molecular mechanisms of ARF (Kaur et al. 2002; Steffens & Rasmussen 2016), including transcriptomic and related omics approaches (Chen et al. 2020; Wang et al. 2022; Yuan et al. 2024), genetic mapping of root-related traits in *Vitis* has so far been confined to structured populations of one or two species. Blois et al. (2023) analyzed a wild *V. berlandieri* population of 211 grafted genotypes and detected 11 QTL for four root-related traits, accounting for up to 25.1% of the variance, and Morel et al. (2026) mapped early root-related traits in a *V. berlandieri* × *V. rupestris* population of 449 genotypes propagated both as hardwood cuttings and as grafted plants, where individual QTL explained 3.1 to 14.1% of the phenotypic variance and the regions detected were largely non-overlapping between the two propagation methods. No study has yet mapped adventitious rooting across a broad, multi-species *Vitis* germplasm collection. Genetic mapping has been used for this trait in other model and crop species. In *Populus trichocarpa*, for example, a GWAS identified 277 unique loci associated with ARF (Nagle et al. 2023), and in rose a GWAS identified a putative phosphoinositide phosphatase for marker-assisted selection (Wamhoff et al. 2023).

In this study, we leveraged the extensive genetic diversity of 308 accessions spanning 18 *Vitis* species, including many underutilized wild taxa, to investigate the genetic basis of adventitious root formation. Accessions were evaluated over three growing seasons, and we additionally derived an index describing the transition from callus-stage to post-transplant rooting, quantified season-to-season repeatability, and tested whether rooting is related to the climate of the collection site. By evaluating rooting traits at two developmental stages and incorporating high-resolution genomic and hyperspectral data, we aimed to uncover patterns of natural variation, identify genomic regions associated with ARF-related phenotypes, and evaluate the potential of genomic and phenomic prediction for selecting rooting traits, representing the first application of such approaches in *Vitis* rootstock germplasm.

## Materials and Methods

### Plant material, experimental design, and phenotyping

A total of 308 genotypes representing 18 *Vitis* species (Table 1) were evaluated using plant material collected over three growing seasons (2024, 2025, and 2026). Most accessions originated from the southwestern United States and are part of the University of California, Davis grapevine germplasm collection, which has previously been used to investigate biotic and abiotic stress tolerance (Heinitz et al. 2020; Morales-Cruz et al. 2021, 2023; Riaz et al. 2018; Riaz et al. 2020; Sharma et al. 2024). The species included in the study were: *V. acerifolia*, *V. aestivalis*, *V. arizonica*, *V. berlandieri*, *V. californica*, *V. candicans*, *V. champinii*, *V. cinerea*, *V.* × *doaniana*, *V. girdiana*, *V. labrusca*, *V. monticola*, *V. riparia*, *V. rupestris*, *V. shuttleworthii*, *V. simpsonii*, *V. treleasei*, and *V. vulpina*.

**Table 1.**
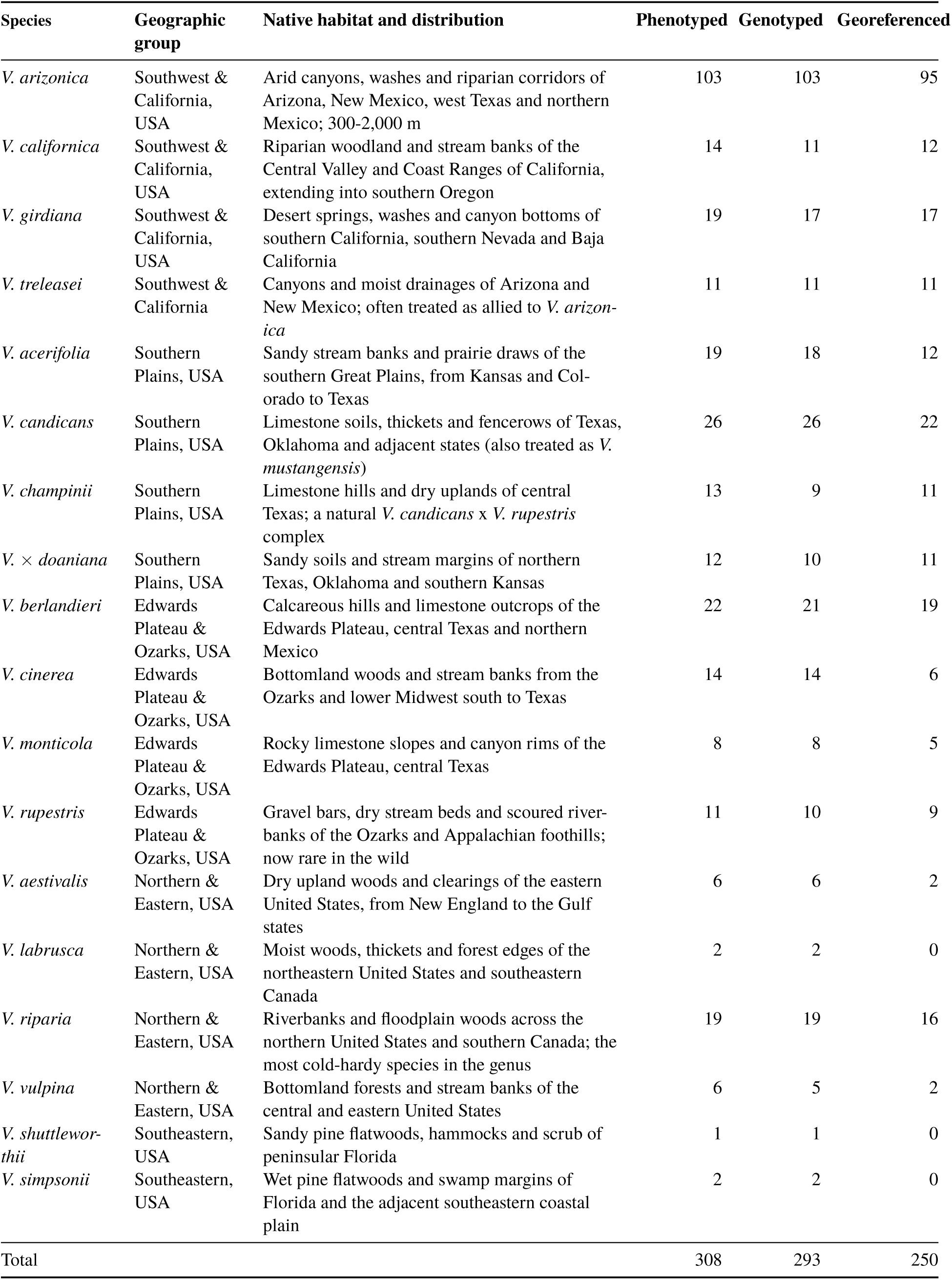
The 18 *Vitis* species represented in the panel, their native habitat and distribution, and the number of accessions of each.

Dormant bud cuttings were collected from field-grown mother vines maintained at UC Davis and propagated and scored following the workflow in Figure 1. Collections were carried out early in the morning (between 7:00-10:00 AM) in January of each year. For each genotype, twelve cuttings were selected, bundled, and soaked overnight in a 0.1% sodium chlorite (NaClO_3_) solution for surface sterilization. The following day, cuttings were transferred to an airtight container with callusing media and incubated at 29 °C for two weeks. After this period, the top bud of each cutting was waxed to minimize dehydration, and the cuttings were planted in 15 × 5 × 5 cm sleeves filled with a 1:1 mixture of callusing media and peat moss. The cuttings were thoroughly watered and returned to the callusing room for an additional five days to promote bud emergence. They were then moved to a growth chamber with bottom heat (27 °C) and maintained for two weeks before being transferred to a greenhouse for an additional six weeks of growth. This protocol mirrors commercial callusing practice, in which cuttings are held warm and moist for two to three weeks before being planted out.

**Figure 1.**
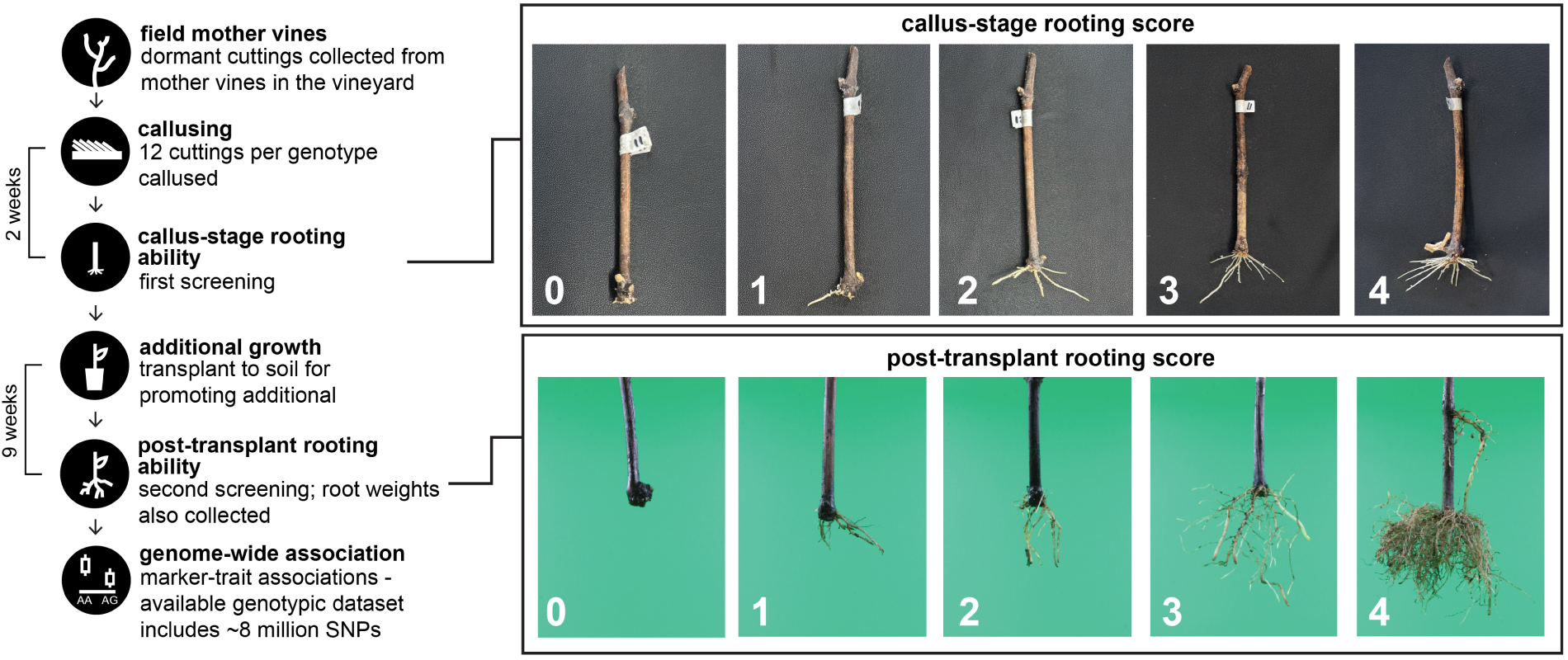
Schematic workflow showing plant material collection, processing, and root scoring at the two developmental stages.

After callusing induction, cuttings were evaluated for callus-stage rooting ability using an ordinal system ranging from 0 (no callus/roots) to 4 (maximum roots; Figure 1). The weight of each cutting was recorded at this point. Cuttings were not standardized for length or diameter, so this weight is an index of how much material each cutting started with rather than a trait selected for in its own right; it was recorded to test whether rooting performance was related to the size of the starting material. After six weeks of growth in the greenhouse, six replicates of each genotype were randomly selected for further evaluation. The soil was carefully removed, roots were washed and cleaned, and post-transplant rooting ability was assessed using the same 0-4 ordinal scale. Root biomass was recorded immediately after post-transplant scoring.

Because of logistical reasons, measurement structure differed among seasons. In 2024, callus-stage rooting was scored once per bundle, so a single integer was recorded per accession; post-transplant scores were taken on each of six plants and averaged, producing fractional values. In 2025, individual cuttings were followed and means were obtained for both stages. In 2026, only an average score per bundle was recorded, and no weights were taken. Consequently, each accession contributes at most one value per season per trait, and root weight and cutting weight are available for 2024 and 2025 only. Supplemental Table S1 summarizes the measurement basis by season and trait.

To describe the transition between the two developmental stages, we defined a transplant-response index as the difference between the within-season standardized post-transplant score and the within-season standardized callus-stage score for the same accession. Standardizing within season before differencing places both stages on a common scale and prevents the index from being dominated by whichever stage happened to have the larger variance in a given year. A positive value indicates an accession that performs better after transplanting than its callus-stage score would predict.

### Statistical analysis

All analyses were performed in R (R Core Team 2024) with mixed models fitted by REML in lme4 (Bates et al. 2015).

Because each accession contributes at most one observation per season, genetic and error variance within a season are not separately identifiable, and a plot-basis broad-sense heritability cannot be estimated from these data. We therefore report repeatability (broad-sense heritability on an entry-mean basis) and state this explicitly wherever a heritability-like quantity is used.

For each trait we fitted

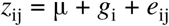

where *z*_ij_ is the within-season standardized value for accession *i* in season *j*, *g*_i_ ∼ N(0, σ^2^_a_) is the random effect of accession, and *e*_ij_ ∼ N(0, σ^2^_e_). Best linear unbiased predictions (BLUPs) of *g*_i_ were used as the phenotype for all downstream analyses. Repeatability was computed three ways: on a single-season basis, σ^2^_a_/(σ^2^_a_ + σ^2^_e_); on an entry-mean basis using the harmonic mean number of seasons per accession, σ^2^_a_/(σ^2^_a_ + σ^2^_e_/*n*); and as the generalized heritability of Cullis et al. (2006), 1 − PEV/σ^2^_a_, which accounts for the unbalanced number of seasons per accession. Confidence intervals were obtained from 1,000 parametric bootstrap replicates. The significance of σ^2^_a_ was tested by likelihood-ratio test against the null of no accession effect, using the 50:50 mixture of 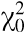and 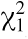 appropriate for a variance component on the boundary.

To separate between-species from within-species genetic variance, we additionally fitted a model with species and accession-within-species as nested random effects and report the proportion of genetic variance attributable to species.

Agreement between seasons for the two ordinal scores was quantified with quadratic-weighted Cohen’s kappa (Cohen 1968) with 1,000 replicate accession bootstrap confidence intervals, computed between seasons for the same accession. Where a season’s values are means rather than native integers, they were rounded to the nearest whole score before the kappa was computed, and the Spearman correlation on the unrounded values is reported alongside (Supplemental Figure S1).

### Accession’s original collection site

Wild collection sites were georeferenced for 250 of the 308 accessions. Bioclimatic variables were extracted from WorldClim 2.1 (Fick & Hijmans 2017) at 2.5 arc-minute resolution using geodata and terra.

For each trait × climate variable combination, we fitted a linear mixed model of the trait BLUP on the standardized climate variable with species as a random intercept, so that only variation within species contributes to the slope. We report both this within-species slope and the plain correlation across all georeferenced accessions; the two answer different questions, because species differ in both where they grow and how well they root, and the plain correlation therefore contains whatever the species composition of the panel contributes. Multiplicity across 85 trait × variable combinations was controlled with the Benjamini–Hochberg procedure (Benjamini & Hochberg 1995).

### Genotyping and genome-wide association

Sequencing data were compiled from previously published studies (Morales-Cruz et al. 2021; Morales-Cruz et al. 2023; Cochetel et al. 2023) and are publicly available in the NCBI Sequence Read Archive under BioProject IDs PRJNA731597, PRJNA842753, and PRJNA984685. Raw reads were filtered and quality-checked using Trimmomatic v0.36 (Bolger et al. 2014) and FastQC v0.11.9 (Andrews 2010), scanned with a sliding window of four base pairs and trimmed when average base quality dropped below Q20; leading and trailing bases below Q3 were removed and only reads ≥60 bp retained. Filtered reads were aligned to haplotype 1 of the *Vitis girdiana* SC2 genome v1.0 (Cochetel et al. 2023) using BWA-MEM (bwa v0.7.12-r1039; Li 2013). Joint genotyping was performed using GATK v4.0.12.0 (Van der Auwera & O’Connor 2020) as described in Cochetel et al. 2023. Briefly, duplicates were marked using Picard (v2.8.2- 4-g2105a1e-SNAPSHOT, https://broadinstitute.github.io/picard/) MarkDuplicates and read groups added with AddOrReplaceReadGroups. Variant calling used HaplotypeCaller (ploidy = 2; minimum base quality Q20) and genotypes were consolidated with GenotypeGVCFs. SNP filtering was conducted with bcftools v1.19 (Danecek et al. 2021) applying GATK hard-filtering thresholds (QD < 2 || FS > 60 || SOR > 3 || MQ < 40 || MQRankSum < -12.5 || ReadPosRankSum < -8); only biallelic SNPs were retained, with a maximum of 25% missing data. Genotypes were available for 293 of the 308 accessions. All accessions were aligned to a single haplotype-resolved assembly so that variants share one coordinate system, which a joint association analysis requires. A super-pangenome assembled from the same panel is available (Cochetel et al. 2023); reads from the species most divergent from *V. girdiana* are expected to align and to be called slightly less completely against a single haplotype, and the consequences of this reference bias are considered in the Discussion.

Genotype data were imported with PLINK v1.9 (Chang et al. 2015) and filtered at a minor allele frequency of 0.05, yielding 3,355,029 SNPs from 4,087,021 called variants. GWAS was conducted using a mixed linear model implemented in GEMMA v0.98.3 (Zhou & Stephens 2012). A standardized genomic relationship matrix was computed with -gk 2 and the first ten principal components, computed in PLINK from the same marker set, were included as fixed covariates. Genome-wide significance was declared at a Bonferroni threshold of α/m with α = 0.05, corresponding to p ≤ 1.49 × 10^−8^ for 3,355,029 tests.

### Candidate genes at associated loci

The size of the interval searched for candidate genes was chosen empirically. Linkage disequilibrium was measured in this panel from 292, 342 marker pairs in a 50,000-SNP subset: mean r^2^ was 0.179 at 2 kb, 0.127 at 8 kb, 0.102 at 25 kb, 0.086 at 50 kb, and 0.078 at 100 kb, against a background of 0.071 for pairs more than 400 kb apart. LD therefore halves within approximately 9 kb and reaches background by roughly 50 kb. We retained a ±100 kb window for the candidate gene search as a conservative net, and report the distance from each gene to the lead SNP, defined as the most significant SNP at a locus after clumping.

Gene models and functional annotations were taken from the *V. girdiana* SC2 v1.0 haplotype 1 annotation (www.grapegenomics.com). Genes overlapping each ±100 kb window were extracted from the GFF3 and joined to the functional annotation. Each gene is reported with its distance to the lead SNP and a flag indicating whether it falls within the ±25 kb core, so that a tighter window can be applied without a second table (Supplemental Table S2).

### Spectral data

Hyperspectral reflectance was collected in the leaves at the time post-transplant rooting scores were recorded, following the protocol of Sharma et al. (2024) and Lupo et al. (2025). Spectra were interpolated to 1 nm resolution over 380-1100 nm, giving 721 wavelengths. A total of 1,408 scans were recorded; eight scans labeled as instrument tests were removed, and the remaining 1,400 scans were screened for missing values, flat traces, and low correlation to the median spectrum of their own sample. Scans were averaged to a single spectrum per accession, giving 235 accessions with a spectrum (median six scans each).

We compared four preprocessing pipelines: raw interpolated reflectance; standard normal variate (SNV) scatter correction (Barnes et al. 1989); and SNV followed by first- and second-derivative Savitzky-Golay filtering (Savitzky & Golay 1964; window 11, polynomial order 2).

### Genomic and phenomic prediction

Prediction used GBLUP,

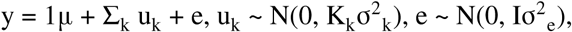

with variance components estimated by REML. Three relationship matrices were constructed: an additive genomic relationship matrix following VanRaden (2008) from nested marker subsets of 5,000, 10,000, 20,000, and 50,000 SNPs, each subset being contained within the next larger one; a spectral relationship matrix computed identically from centered and scaled reflectance; and a taxonomy matrix in which accessions of the same species take the value 1 and 0 otherwise. A combined genomic-plus-spectral model was fitted as a two-kernel model so that the variance attributable to each source is separately estimable.

Five validation scenarios were evaluated: within-season prediction for 2024, 2025, and 2026, and two across-season designs training on 2024 and validating on accessions evaluated only in 2025 or only in 2026, so that validation genotypes are genuinely unseen. Within-season accuracy was assessed by five-fold cross-validation replicated 50 times; accuracy is reported as the Pearson correlation between observed and predicted values pooled across folds, together with the regression slope of observed on predicted as a measure of calibration.

The comparison among genomic, spectral, taxonomy, and combined models used only the 235 accessions carrying phenotypes, markers, and spectra, so that model differences are not confounded with differences in sample composition.

## Results

### Phenotypic variation in adventitious rooting across 18 Vitis species

We evaluated adventitious root formation in 308 accessions spanning 18 *Vitis* species over three consecutive seasons. For callus-stage rooting, 154 accessions were evaluated in all three seasons (Figure 2A), 147 in two, and 7 in one; for post-transplant rooting, the corresponding numbers were 150, 138, and 20. Root weight and cutting weight were recorded in 2024 and 2025 only, with 155 and 159 accessions respectively evaluated in both seasons (Figure 2A). Trait values differed among seasons in both location and spread, most conspicuously for the post-transplant score in 2026 (Figure 2B).

**Figure 2.**
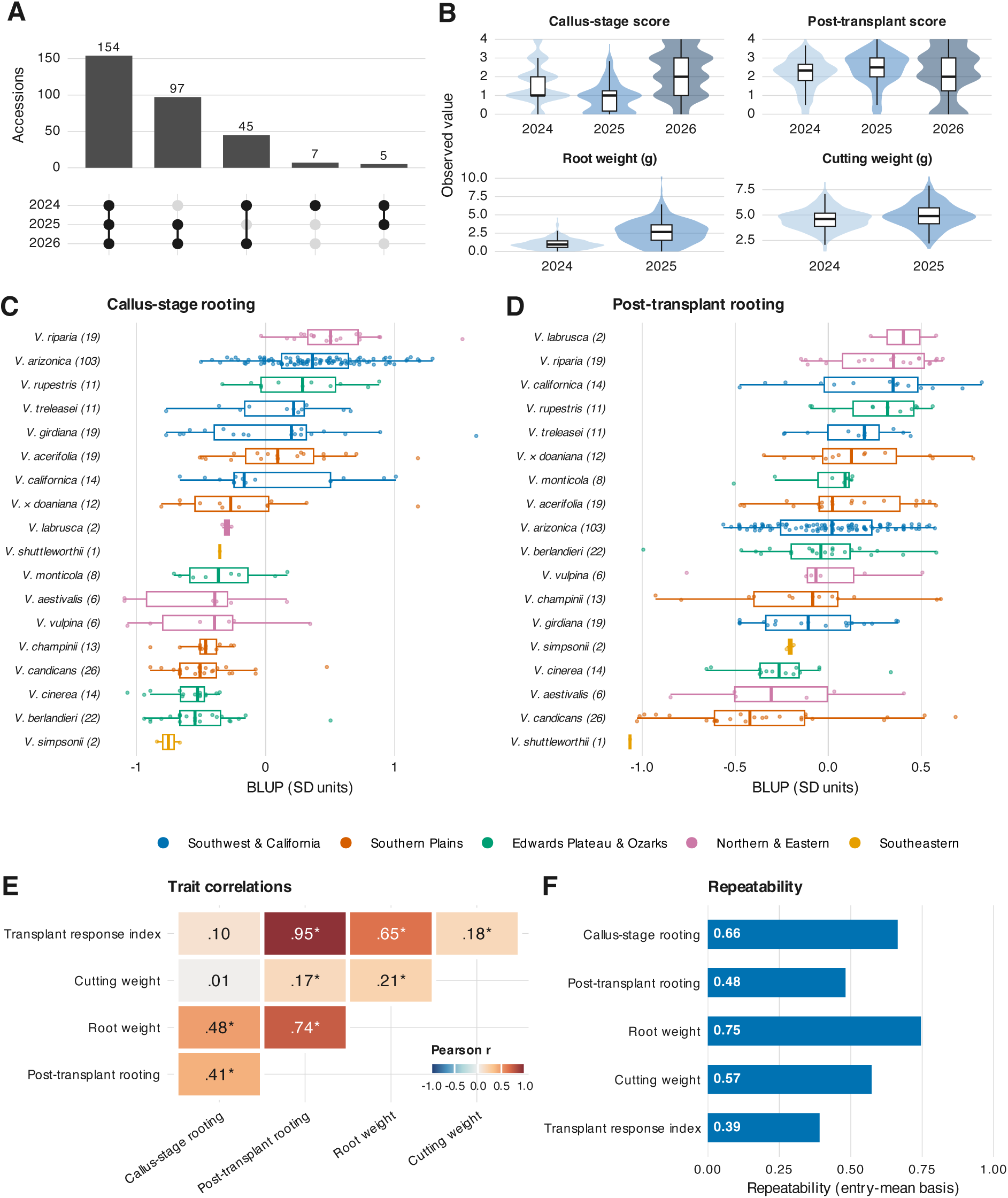
Phenotypic variation in adventitious root formation across 308 accessions and 18 *Vitis* species. (A) Number of accessions evaluated in one, two, or three seasons, by trait. (B) Distribution of each trait by season. (C, D) Trait BLUPs by species. (E) Pairwise correlations among trait BLUPs; asterisks mark p < 0.05. (F) Repeatability on an entry-mean basis for each trait; root weight and cutting weight use two seasons (2024-2025), the remaining traits three.

Callus-stage rooting BLUPs ranged from 0.50 (T52) to 3.22 (SC42, *V. girdiana*), with a panel mean of 1.62. Species with the highest mean callus-stage rooting were *V. riparia* (2.12), *V. arizonica* (1.98), and *V. rupestris* (1.86); the lowest were *V. simpsonii* (0.84), *V. cinerea* (0.98), and *V. aestivalis* (1.08) (Figure 2C). The top-performing accessions were SC42 (*V. girdiana*, 3.22), OK14-027 (*V. riparia*, 3.11), UT12-064 (*V. arizonica*, 2.88), AZ11-100 (*V. arizonica*, 2.78), and NM11-021 (*V. arizonica*, 2.77).

Post-transplant rooting BLUPs ranged from 1.18 (Haines City) to 3.09 (NC9, *V. californica*), with a mean of 2.25. The highest species means were *V. labrusca* (2.67), *V. riparia* (2.55), and *V. rupestris* (2.53); the lowest were *V. shuttleworthii* (1.18), *V. candicans* (1.89), and *V. cinerea* (2.00) (Figure 2D). The best-performing accessions were NC9 (*V. californica*, 3.09), OK12-024 (*V.* × *doaniana*, 3.05), NC5 (*V. californica*, 2.99), OK12-019 (*V. candicans*, 2.95), and NM12-114 (*V. riparia*, 2.88).

Root weight BLUPs ranged from 0.44 (Haines City) to 5.82 (C78-95, *V. californica*), with a mean of 1.89. The highest species means were *V. californica* (3.05), *V. labrusca* (2.77), and *V. rupestris* (2.61); the lowest were *V. shuttleworthii* (0.44), *V. candicans* (1.04), and *V. cinerea* (1.15). The greatest root weight was produced by C78-95 (*V. californica*, 5.82), SC42 (*V. girdiana*, 5.57), NC55 (*V. californica*, 4.89), Vru 94 (*V. rupestris*, 4.66), and NC5 (*V. californica*, 4.47). Species rankings were therefore trait-dependent. *V. riparia*, *V. rupestris* and *V. californica* were in the top five for every trait, and *V. cinerea* and *V. candicans* in the bottom five for root weight and post-transplant rooting, but *V. arizonica* ranked second at the callus stage and eighth after transplanting, and *V. berlandieri* fifteenth of eighteen at the callus stage and eleventh after transplanting.

### Trait correlations, repeatability, and the structure of genetic variance

Cutting weight, an index of the size of the starting material, showed no correlation with callus-stage rooting (r = 0.01, p = 0.92) and correlations that were statistically significant but weak with post-transplant rooting (r = 0.17, p = 0.003) and root weight (r = 0.27, p ≤ 0.001), fewer than 8% of the variance being shared in either case, so how much material a cutting began with explains little of the variation in rooting (Figure 2E). Callus-stage and post-transplant rooting were moderately correlated (r = 0.41, p = 1.1 × 10^−13^), and post-transplant rooting was strongly correlated with root weight (r = 0.74, p = 1.1 × 10^−53^).

Accession effects were highly significant for every trait (likelihood-ratio test against the boundary null; p ≤ 4.3 × 10^−7^ in all cases). Repeatability on an entry-mean basis was highest for root weight (0.74; 95% CI 0.66-0.80) and callus-stage rooting (0.66; 0.58-0.72), intermediate for cutting weight (0.56; 0.43-0.66), and lowest for post-transplant rooting (0.47; 0.35-0.56) and the transplant-response index (0.38; 0.24-0.48) (Figure 2F).

Excluding the 2026 season raised repeatability substantially for post-transplant rooting (from 0.47 to 0.67) and for the transplant-response index (0.38 to 0.56), while leaving callus-stage rooting and the weights essentially unchanged. The 2026 season was scored once per bundle rather than per plant, and post-transplant scores from that season correlate with 2024 at r = 0.16 and with 2025 at r = 0.25, against r = 0.60 between 2024 and 2025. Weighted-kappa agreement between seasons was moderate for callus-stage 2024 vs 2026 (κ_w_ = 0.52, 0.43-0.61) and post-transplant 2024 vs 2025 (κ_w_ = 0.54, 0.41-0.63), but low for any comparison involving the 2026 post-transplant score (κ_w_ = 0.13-0.16) (Supplemental Figure S1).

Partitioning genetic variance between and within species revealed a striking contrast among traits. Between-species differences accounted for 67.9% of the genetic variance in callus-stage rooting but only 49.8% in post-transplant rooting, 42.1% in root weight, 38.7% in the transplant-response index, and 9.6% in cutting weight. Early-stage rooting is therefore substantially more phylogenetically structured than the later stages, a pattern that recurs in the prediction results below.

### Rooting is associated with the climate of the collection site

Wild collection sites were available for 250 accessions across 15 species, spanning a wide range of precipitation and temperature regimes across western and central North America (Figure 3A, B). Of 85 trait × climate combinations tested, 7 reached a Benjamini–Hochberg q < 0.05 and 12 reached q < 0.10 (Figure 3C, Supplemental Table S3). The strongest association was between root weight and precipitation of the driest month (within-species standardized slope β = −0.35, p = 7.9 × 10^−6^, q = 0.0007), followed by callus-stage rooting and the same variable (β = −0.19, p = 1.1 × 10^−4^) and post-transplant rooting and the same variable (β = −0.15, p = 1.4 × 10^−4^). Associations with precipitation of the driest quarter, precipitation of the warmest quarter, and annual precipitation were of the same sign and comparable magnitude (Figure 3D).

**Figure 3.**
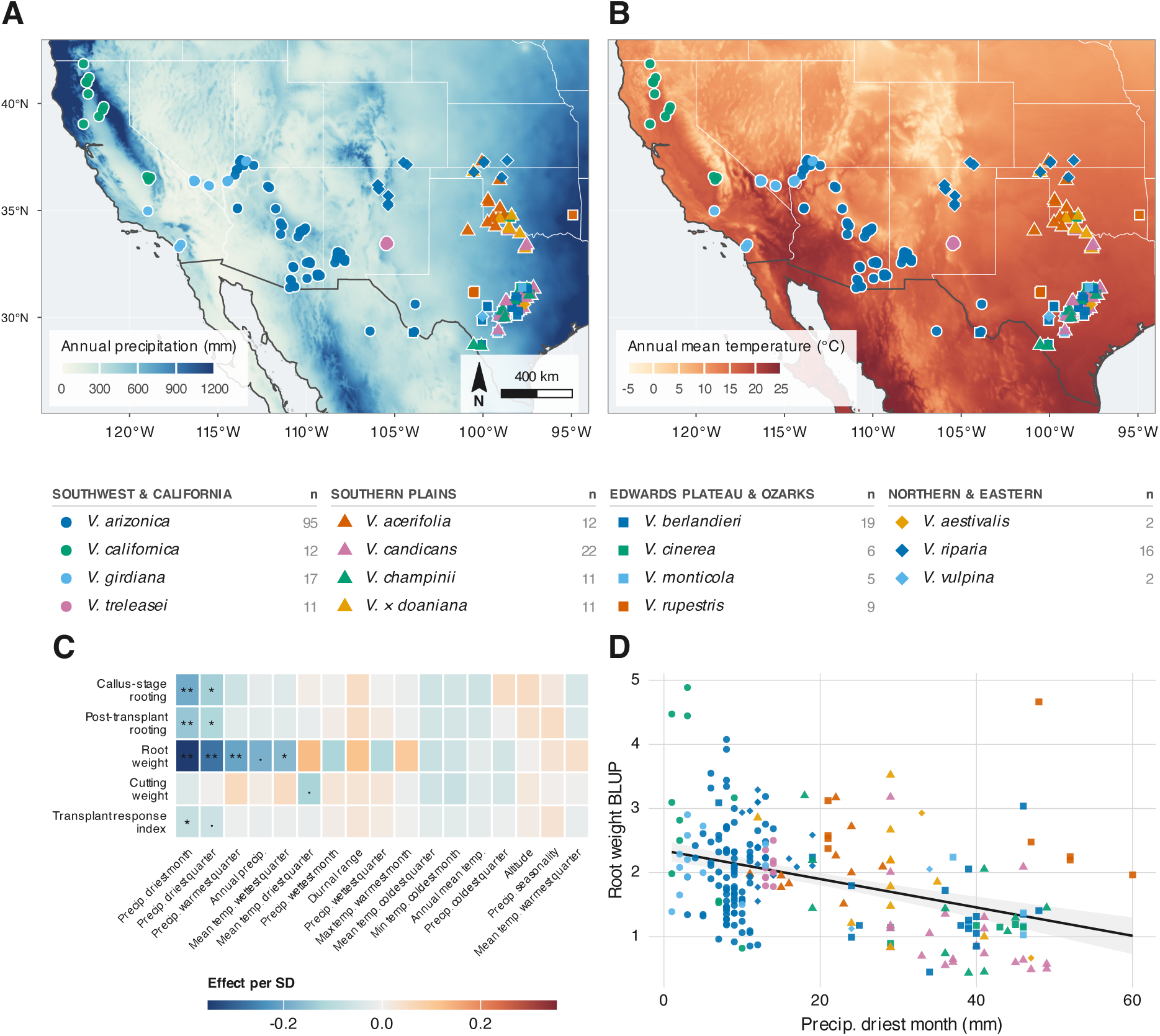
Geographic distribution, climatic context, and climate associations of the *Vitis* accession panel. (A, B) Collection sites of 250 of the 308 phenotyped accessions, representing 15 species, plotted over annual precipitation and annual mean temperature. Point shape denotes the broad geographic origin of the species and point color denotes species within each origin. (C) Association between the climate of the collection site and each rooting trait, expressed as the change in trait BLUP per standard deviation of the climate variable. Asterisks mark Benjamini–Hochberg q < 0.01 () and q < 0.05 (*); a mid-dot marks q < 0.10. (D) The strongest of these associations, root weight against precipitation of the driest month, with a linear fit and 95% confidence band

The direction is consistent across traits and survives the removal of species means; accessions collected from sites with lower dry-season precipitation root better and produce more root biomass. Because species differ in both their geographic ranges and their rooting ability, the plain across-panel correlations are larger than the within-species slopes (for example r = −0.54 versus β = −0.19 for callus-stage rooting and precipitation of the driest month), and both are reported. We emphasize that these are observational associations and do not establish causation.

### GWAS revealed loci and candidate genes associated with rooting

Association analysis was performed on over 3.3 million SNPs in 293 accessions (284 for root weight, which was not recorded for every accession). Genomic control factors ranged from 0.90 to 1.08 (callus-stage rooting 0.90, root weight 1.01, cutting weight 1.00, transplant response 1.07, post-transplant rooting 1.08), indicating that the kinship matrix and ten principal components adequately controlled population stratification, with no evidence of inflation (Figure 4, Supplemental Table S6). The proportion of phenotypic variance explained by all markers jointly, estimated from the GEMMA null model, was high for every rooting trait: 0.88 ± 0.05 for root weight, 0.84 ± 0.09 for callus-stage rooting, 0.77 ± 0.11 for post-transplant rooting and 0.76 ± 0.12 for the transplant-response index, against 0.54 ± 0.22 for cutting weight. These values are not narrow-sense heritabilities in the usual sense; in a panel spanning 18 species the genomic relationship matrix captures both within- and between-species relatedness, however, they confirm that the traits are strongly structured by genotype.

**Figure 4.**
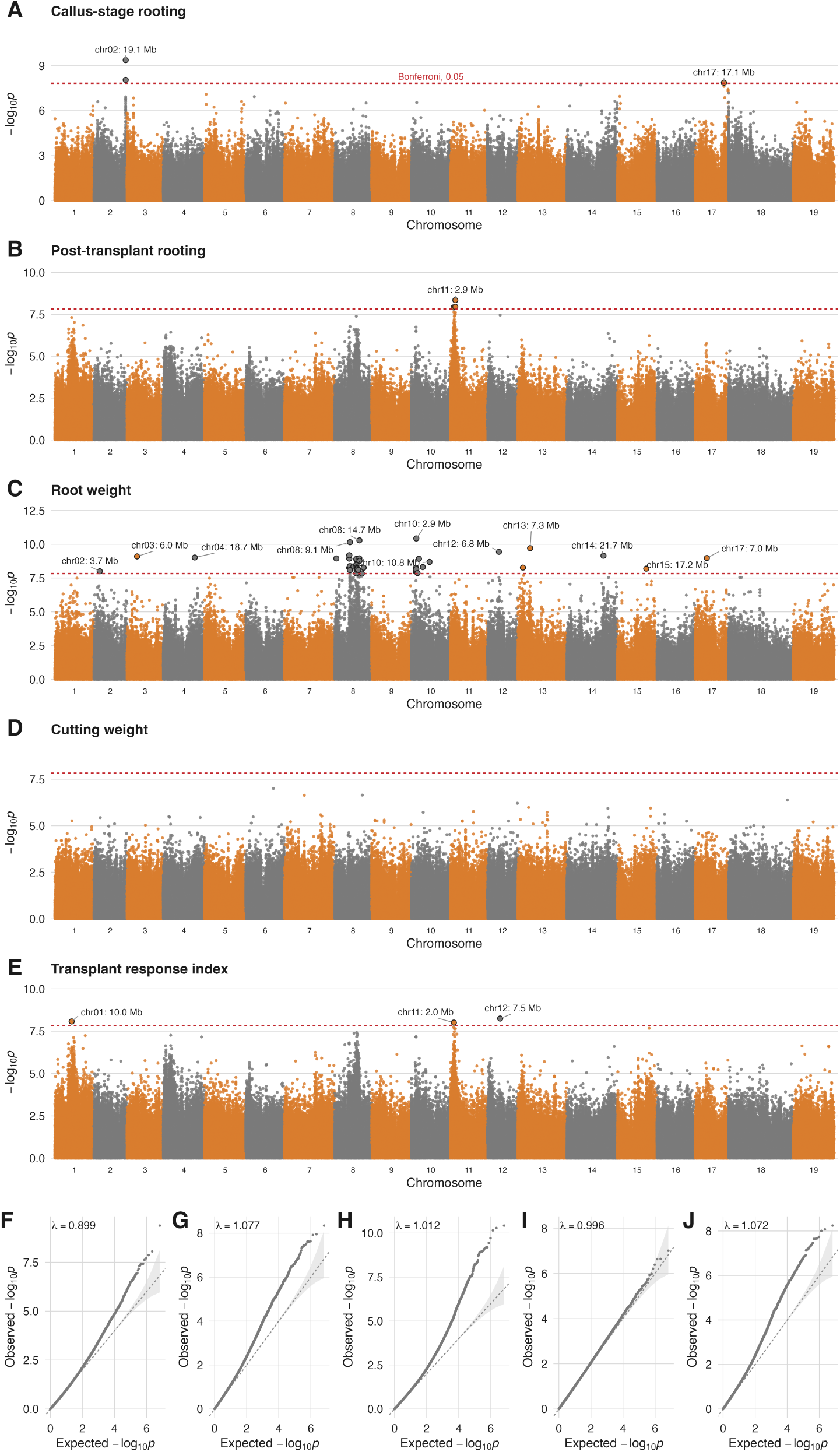
Genome-wide association results for the five rooting traits. Manhattan plots of −log_10_(p) from a mixed linear model in GEMMA fitting a standardized kinship matrix and ten principal components, with a narrow quantile–quantile plot beside each. The dashed red line is the Bonferroni threshold at α = 0.05. SNPs passing it are drawn with a dark outline in the color of their chromosome. Faint vertical rules mark lead positions across all panels to aid comparison between traits.

Fifty-four SNPs exceeded the Bonferroni threshold across the four rooting traits: three for callus-stage rooting, three for post-transplant rooting, forty-five for root weight, and three for the transplant-response index. These resolved into 18 independent loci: two for callus-stage rooting (chromosomes 2 and 17), one for post-transplant rooting (chromosome 11), twelve for root weight (chromosomes 2, 3, 4, 8 [two loci], 10 [two loci], 12, 13, 14, 15 and 17), and three for the transplant-response index (chromosomes 1, 11 and 12). Root weight, the trait with the highest repeatability and the highest marker-based variance component, therefore accounts for two thirds of the detected loci; the two rooting scores, which are recorded on a coarse ordinal scale, each yielded a single well-resolved region plus one secondary signal (Figure 4).

Effect sizes are reported on the scale of the trait BLUPs (Table 2). The strongest single association was on chromosome 10 at 2,946,219 bp for root weight (*p* = 3.7 × 10^−11^, β = 1.205 ± 0.175, minor allele frequency 0.118), followed by two loci on chromosome 8 at 14,735,345 bp (*p* = 5.1 × 10^−11^, β = 1.410 ± 0.206) and 9,066,599 bp (*p* = 7.0 × 10^−11^, β = 1.446 ± 0.213), both at a frequency of 0.117. For callus-stage rooting the lead SNP on chromosome 2 at 19,087,797 bp (*p* = 4.2 × 10^−10^, β = 0.374 ± 0.058) is the only common-frequency lead in the set (0.463); every other lead has a minor allele frequency between 0.049 and 0.334, which is expected in a wild panel where favorable alleles are often confined to one or a few species. Post-transplant rooting was associated with a single locus on chromosome 11 at 2,905,633 bp (*p* = 4.5 × 10^−9^, β = −0.434 ± 0.072, frequency 0.334), and the transplant-response index with loci on chromosome 12 at 7,536,289 bp, chromosome 1 at 9,992,637 bp and chromosome 11 at 2,039,493 bp.

**Table 2.**
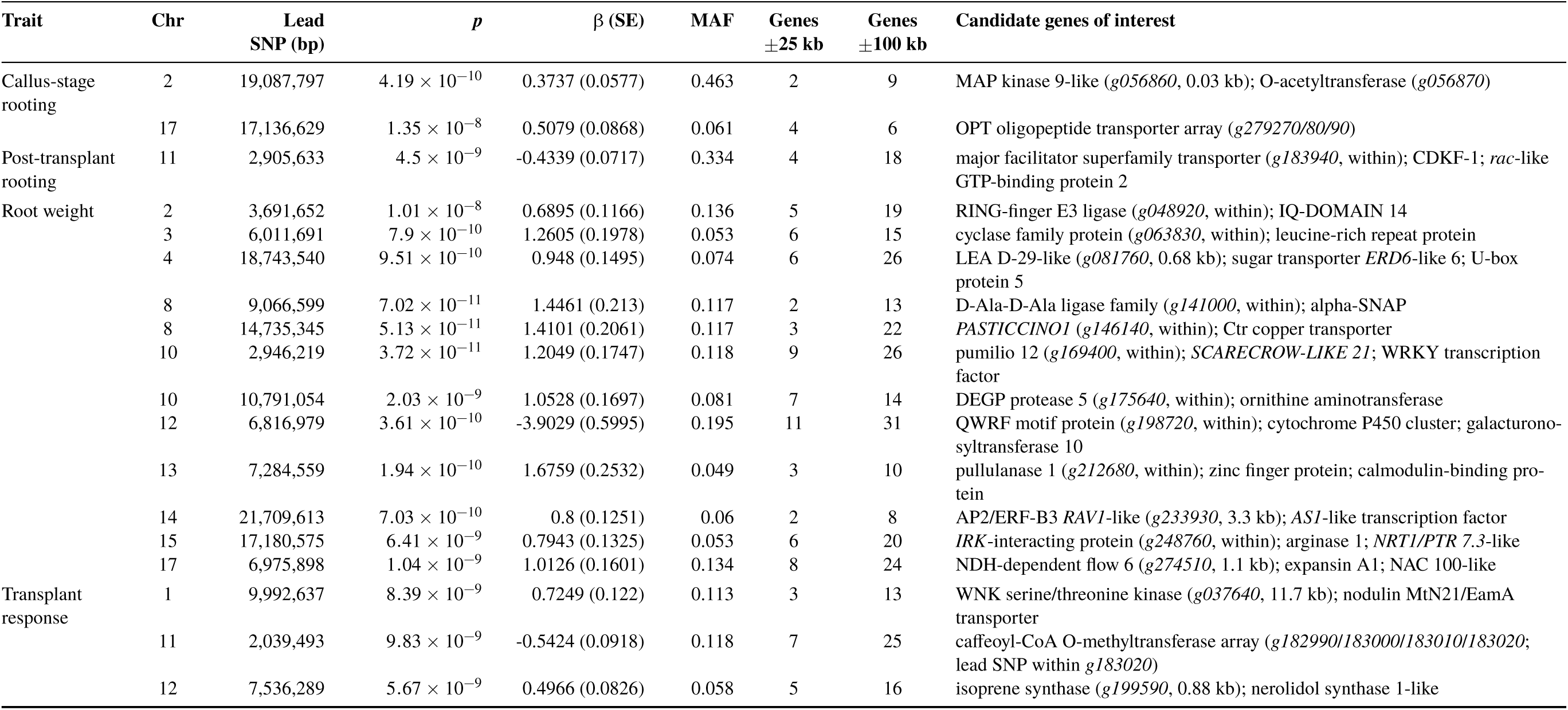
The 18 loci significantly associated with rooting traits. For each locus: trait, chromosome, position of the lead SNP on haplotype 1 of the *V. girdiana* SC2 assembly, Wald *p*-value, effect size β and its standard error on the scale of the trait BLUPs, minor allele frequency, the number of annotated genes within the ±25 kb linkage-disequilibrium core and within the full ±100 kb search window, and the genes of most interest in the core with their distance from the lead SNP in parentheses.

Each locus is summarized in Table 2, and every gene within ±100 kb of a lead SNP is reported with its distance and orientation and a flag marking the ±25 kb linkage-disequilibrium core (Supplemental Table S2). Ninety-three of the 315 genes fall within that core, an average of five per locus. At every locus the association signal is confined to a narrow interval containing only a handful of annotated genes (Figure 5).

**Figure 5.**
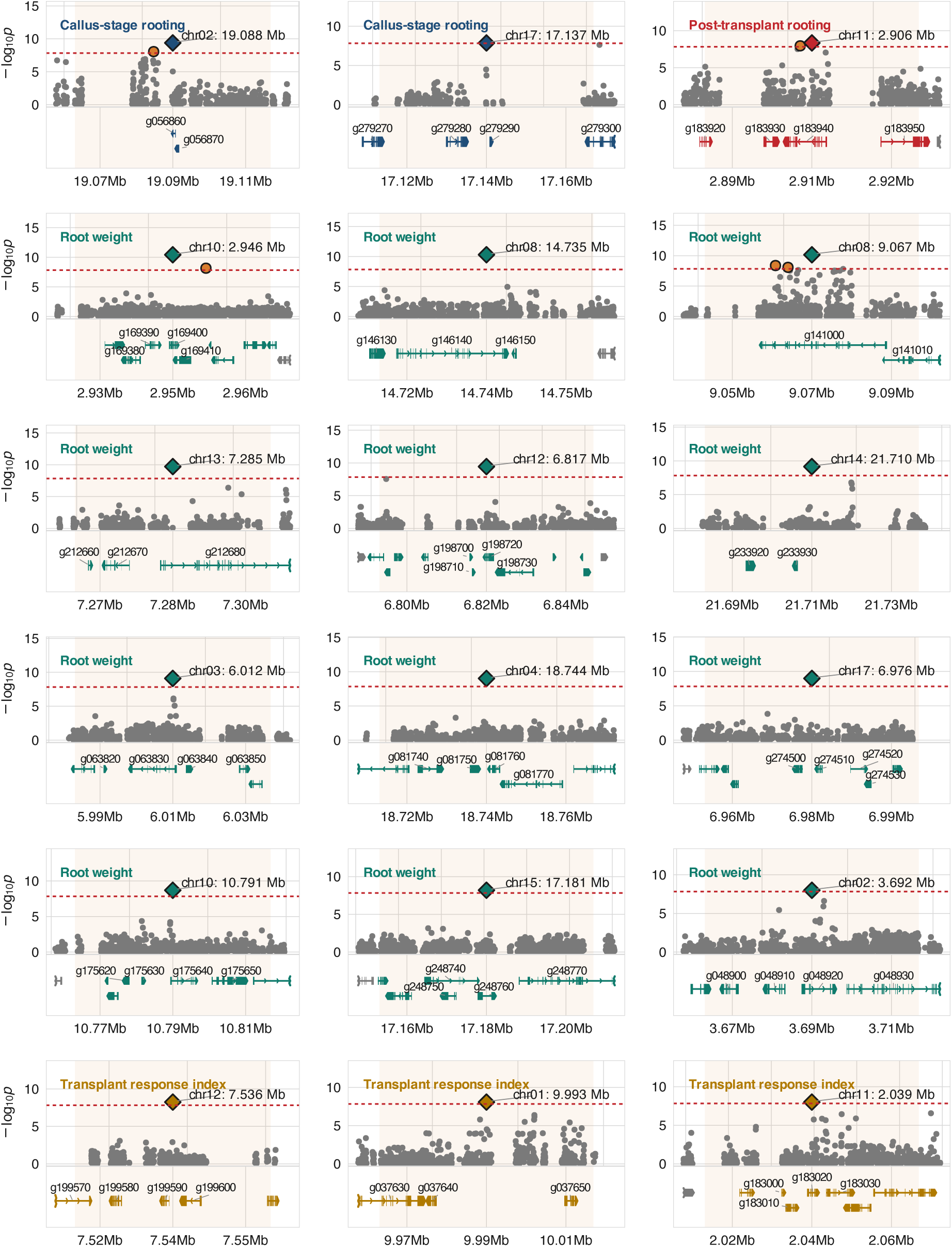
Local views of the 18 significant loci. For each QTL, the upper panel shows −log_10_(p) across a ±30 kb window centered on the lead SNP, with the lead SNP marked by a diamond, and the lower panel shows the gene models in the same interval each drawn as exon boxes on an intron line. The shaded band marks the ±25 kb interval within which LD remains appreciable in this panel. Gene models are clipped to the plotted interval, so exon structure lying outside the window is not shown.

Several of these are directly interpretable in the context of adventitious rooting. The callus-stage locus on chromosome 2 places the lead SNP 30 bp from *g056860*, a mitogen-activated protein kinase 9-like gene, with an O-acetyltransferase family protein 0.9 kb further on; MAPK cascades transduce the wound and auxin signals that initiate root primordia, and xyloglucan O-acetyltransferases modify cell wall extensibility (Zhong et al. 2020), both consistent with a locus acting at the callus stage. The second callus-stage locus, on chromosome 17, sits inside a tandem array of three OPT family oligopeptide transporters spanning 30 kb.

For root weight, the chromosome 8 locus at 14.74 Mb contains the lead SNP within *g146140*, the grapevine orthologue of *PASTICCINO1*, an FKBP-type peptidyl-prolyl isomerase required for auxin-dependent control of cell proliferation and a well-characterized regulator of lateral and adventitious root initiation; this is the most compelling single candidate in the set. The chromosome 10 locus at 2.95 Mb contains *SCARECROW-LIKE 21* (a GRAS-family transcription factor of the SHORT-ROOT/SCARECROW module that specifies root radial patterning) 12.7 kb upstream, together with a WRKY transcription factor 8.1 kb upstream and *pumilio 12* containing the lead SNP. The chromosome 14 locus is flanked by two transcription factors, an AP2/ERF-B3 *RAV1*-like gene 3.3 kb upstream and an *ASYMMETRIC LEAVES 1*-like gene 13.6 kb upstream; AP2/ERF factors integrate age and wound signals during root regeneration (Ye et al. 2020). The chromosome 17 locus at 6.98 Mb includes *expansin A1* 17.1 kb upstream and a *NAC domain-containing protein 100*-like gene 18.9 kb downstream, both cell wall and differentiation associated. The chromosome 15 locus contains an *IRK*-interacting protein at the lead SNP, with *arginase 1* and a *NRT1/PTR FAMILY 7.3*-like nitrate transporter (Wang et al. 2009) within 25 kb. The chromosome 12 root weight locus resolves to a cluster of three cytochrome P450 genes and a probable galacturonosyltransferase 10 within 4 kb of the lead SNP, and the chromosome 4 locus to a late-embryogenesis-abundant protein, a zinc finger, and a sugar transporter *ERD6*-like 6, which is relevant given that adventitious rooting in hardwood cuttings draws on stored carbohydrate.

The transplant-response index gave the most compositionally distinctive locus of the study. On chromosome 11 at 2.04 Mb the lead SNP lies inside *g183020*, one of four caffeoyl-CoA O-methyltransferase genes arrayed within 18 kb of each other. CCoAOMT catalyzes a committed step of monolignol biosynthesis, and lignification and suberization of the stem base are central to how a rooted cutting survives transplanting; a tandem array of these genes at the strongest locus for a trait defined as the change in rooting score between the callus stage and post-transplant is a coherent result rather than a coincidence. The chromosome 1 locus for the same trait contains a nodulin MtN21/EamA-like transporter 18.2 kb downstream (the family that includes the UMAMIT auxin efflux carriers) and a WNK-family serine/threonine kinase 11.7 kb upstream. The chromosome 12 locus for this trait sits between two isoprene synthase genes and a nerolidol synthase, a terpene synthase cluster whose connection to rooting is not obvious and which we report without interpretation.

The single locus for post-transplant rooting, on chromosome 11 at 2.91 Mb, has the lead SNP within a Major Facilitator Superfamily transporter, with a cyclin-dependent kinase F-1 8.0 kb upstream and a *rac*- like GTP-binding protein 2, a ROP GTPase of the type that controls polar cell expansion and root hair initiation, 23.7 kb upstream.

Cutting weight yielded no significant associations at any locus. This measurement was recorded to test whether the size of the starting material was related to rooting, not as a rooting trait in its own right, and cuttings were not standardized for length or diameter, so a flat association profile carries no implication for the rooting results. Its moderate repeatability (0.56) does indicate consistent differences among accessions in the size of the canes they produce, but only 9.6% of that variance lies between species and the panel has correspondingly less power for it; a larger or more standardized sample could well detect associations.

### Genomic and phenomic prediction

Analysis of the spectral dataset revealed two major correlation blocks, one spanning approximately 385-690 nm and a second spanning approximately 700-1080 nm (Figure 6A). Correlations between reflectance and trait BLUPs were low to moderate and negative for the rooting traits, strongest between approximately 420- 500 nm and near 680 nm, and near zero across the near-infrared range; cutting weight showed the opposite sign (Figure 6B). Genetic differences accounted for up to 73% of spectral variance among accessions and up to 58% among species, in both cases peaking near 670-680 nm (Figure 6C), indicating that spectral variation carries a strong genetically driven component.

**Figure 6.**
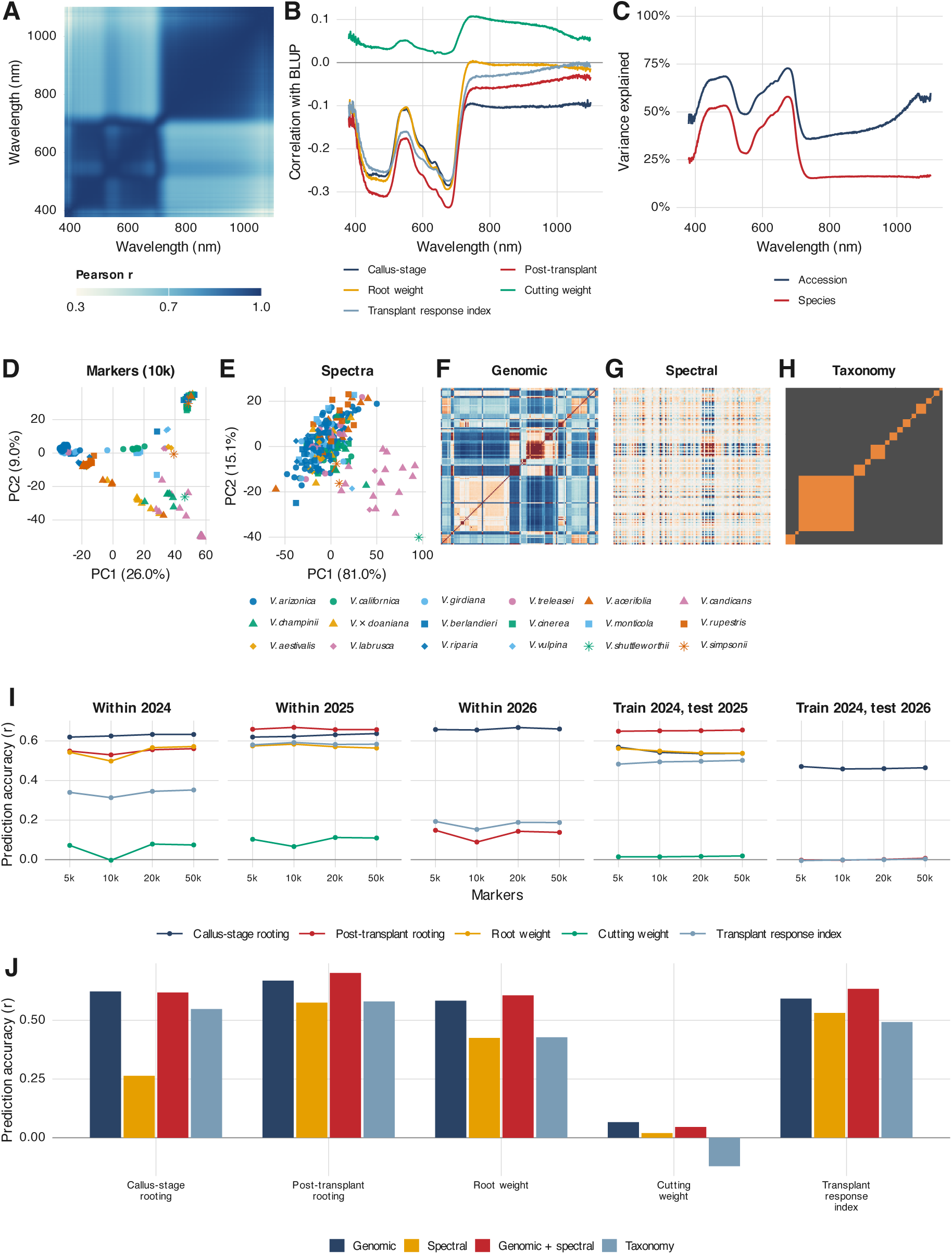
Spectral structure and predictive modeling of adventitious rooting. (A) Correlation between wavelengths across 380- 1100 nm. (B) Correlation between reflectance at each wavelength and each trait BLUP. (C) Proportion of spectral variance explained by differences among accessions and among species. (D, E) Principal component analysis of 283 accessions using 10,000 markers and of 235 accessions using reflectance. (F–H) Genomic, spectral, and taxonomy relationship matrices, accessions ordered by species. (I) Genomic prediction accuracy against marker density for each trait and validation scenario. (J) Genomic, spectral, combined, and taxonomy-based models compared within 2025 at 10,000 markers on the 235 accessions carrying both markers and spectra.

Principal component analysis of the marker data showed pronounced population structure with clear clustering by species, with PC1 and PC2 explaining 26.0% and 9.0% of the variance respectively (Figure 6D). Principal component analysis of the spectral data was dominated by a single component (PC1 81.0%, PC2 15.1%) and showed more diffuse clustering, although some species remained separable (Figure 6E). The genomic, spectral, and taxonomy relationship matrices displayed broadly similar block structure (Figure 6F–H).

Prediction accuracies varied across marker densities and validation scenarios (Figure 6I). Within-season accuracy was highest for callus-stage rooting in 2026 (r = 0.66), post-transplant rooting in 2025 (r = 0.67), and callus-stage rooting in 2024 and 2025 (r = 0.63 in both). Root weight reached r = 0.58 and the transplant- response index r = 0.59, both in 2025. Predicting accessions evaluated only in 2025 from a model trained on 2024 gave r = 0.65 for post-transplant rooting, 0.55 for root weight, 0.54 for callus-stage rooting, and 0.49 for the transplant-response index. Accuracy for genuinely unseen accessions was therefore comparable to within-season accuracy.

Marker density had almost no effect. For callus-stage rooting within 2025, accuracy was 0.620 at 5,000 markers and 0.639 at 50,000, a gain of 0.019 for a tenfold increase in marker number. The same pattern held for every trait and scenario.

Comparing model types within 2025 on the 235 accessions carrying both markers and spectra, the combined genomic-plus-spectral model gave the highest accuracy for post-transplant rooting (0.701 versus 0.668 for genomic alone), root weight (0.605 versus 0.580), and the transplant-response index (0.634 versus 0.592) (Figure 6J). For callus-stage rooting the genomic model was marginally better than the combined model (0.623 versus 0.618), and the taxonomy-only model reached 0.551 for this trait, far above the spectral model at 0.266. For post-transplant rooting and root weight the ordering was reversed, with the spectral model matching or exceeding taxonomy (0.575 versus 0.580, and 0.425 versus 0.427). This mirrors the variance partition: callus-stage rooting is the most phylogenetically structured trait, so species identity alone predicts it well, whereas the later stages carry more within-species variation that spectral data can capture but taxonomy cannot.

Spectral preprocessing effects were trait-dependent. SNV scatter correction followed by second-derivative Savitzky-Golay filtering raised callus-stage accuracy from 0.263 to 0.377, but reduced accuracy for post-transplant rooting (0.575 to 0.568), root weight (0.425 to 0.389), and the transplant-response index (0.531 to 0.491). No single pipeline was best for every trait, so raw reflectance is reported as the primary result, with the full comparison in Supplemental Table S4.

Accuracy as a function of training-set size was close to flat between 50 and 200 accessions for most traits (for example post-transplant rooting: 0.64 at n = 50 and 0.66 at n = 200), indicating that this panel is already near the plateau for these traits (Supplemental Table S5).

Cutting weight was essentially unpredictable in every scenario and by every model (r ≤ 0.17), which is unsurprising for a measurement that indexes the size of the starting material rather than rooting itself and that carries almost no between-species variance. And post-transplant rooting in 2026 was predicted at r = 0.08 within season and r ≈ 0.00 across seasons, against r = 0.67 and 0.65 for the same trait in 2025.

## Discussion

Adventitious root formation in dormant cuttings is the step that determines whether a genotype can be propagated at all. The variation observed in this work across 308 accessions of 18 Vitis species shows that this step is under substantial genetic control while remaining sensitive to the conditions under which cuttings are collected and callused. Rooting in grapevine cuttings responds to temperature, light, humidity and substrate moisture as well as to endogenous hormone levels, carbohydrate reserves, nutrient transport and cutting maturity (Bellini et al. 2014; Da Costa et al. 2013; Druege et al. 2019; Gonin et al. 2019; Pan et al. 2002; Rasmussen et al. 2017; Steffens & Rasmussen 2016). In woody species it originates from preformed or induced primordia associated with dormant buds (Smart et al. 2002). Because commercial propagation depends on this step, small differences in rooting percentage translate directly into nursery throughput (Waite et al. 2015). Against that background, accession effects were significant for every trait and repeatability reached 0.74 for root weight and 0.66 for callus-stage rooting, which is high enough for selection to be worthwhile in a propagation context. The only previous quantitative-genetic estimate for this trait in grapevine, a narrow-sense heritability of 0.31 ± 0.05 for rootstrike across 27 families and 552 individuals (Smith et al. 2013), is lower than the values reported here, as expected given that repeatability also contains non-additive genetic variance and that our estimates are on an entry-mean rather than a single-observation basis.

Many of the species evaluated here are already known as sources of adaptive traits: salinity tolerance (Heinitz et al. 2020; Sharma et al. 2025), drought tolerance (Heinitz et al. 2014; Morales-Cruz et al. 2021) and resistance to pests and diseases (Morales-Cruz et al. 2023; Riaz et al. 2018). Their rooting behavior, however, has not been characterized, which has limited their use as rootstocks and their inclusion in breeding programs. Several non-traditional species rooted as well as or better than the species that dominate commercial rootstock pedigrees. *V. californica* produced the greatest root biomass of any species (mean BLUP 3.05), and *V. girdiana* and *V. treleasei* were competitive with *V. riparia* and *V. rupestris* at the callus stage. The converse also held: not every accession of *V. riparia*, *V. rupestris* or *V. berlandieri* rooted well, and species means concealed wide within-species ranges. Selection for rooting therefore has to be made at the accession level, and species identity is a starting point rather than a substitute for evaluation. Accessions such as SC42 (*V. girdiana*), OK14-027 (*V. riparia*), C78-95 and NC9 (*V. californica*) combined high scores at both stages with high root biomass and are the most immediate candidates for use as rootstocks in their own right or as parents in crosses with elite material.

Scoring rooting at two stages separated by transplanting made it possible to ask whether early and late rooting are the same trait. They are related but not equivalent. Callus-stage and post-transplant scores correlated at r = 0.41, post-transplant rooting correlated with root weight at r = 0.74, and cutting weight was essentially uncorrelated with rooting at either stage, indicating that the size of the starting material has little bearing on whether roots form. The more informative difference between the two stages is not the correlation but the way genetic variance is organized: between-species differences accounted for 68% of the genetic variance at the callus stage but only 50% after transplanting and 42% for root biomass. Early rooting is thus strongly phylogenetically structured, while later performance depends more on variation segregating within species. This has a direct practical consequence, and it is visible again in the prediction results: species identity alone predicts callus-stage rooting reasonably well but cannot rank accessions within a species. Some accessions also showed a mismatch between visual score and biomass, which is expected if genotypes differ in root morphology; Tandonnet et al. (2018) reported that root number tracks fine roots while biomass is driven by coarse roots. More than one metric is needed to describe rooting capacity.

The genetic architecture implied by the association analysis is polygenic, and the loci we detected are its upper tail. Markers jointly explained 76-88% of the variance in the four rooting traits, yet only 18 loci exceeded a genome-wide threshold and each accounted for a similar and modest fraction of residual variance. With 293 accessions, only the largest effects are resolvable, and the loci reported here should be read as the subset that survived a conservative threshold rather than as a complete account. Root weight contributed twelve of the eighteen loci. That is partly biological but mostly methodological: root weight is the only continuous trait in the set and the most repeatable, whereas the two rooting scores are recorded on a 1-5 ordinal scale and yielded three loci between them. The clearest recommendation to come out of this study for future phenotyping is therefore to record a continuous proxy, such as root biomass or an image-derived root area, alongside the visual score. The one comparable genetic dissection of root traits in grapevine reached a similar conclusion from a different direction: in a V. riparia x ’Seyval blanc’ F2 population, 31 QTL were detected for 11 root system architecture traits, with predictive models explaining 13 to 32% of phenotypic variation (Alahakoon & Fennell 2023). Neither that study nor ours resolves a locus of large effect. Loci were detected on several of the same chromosomes in both studies (1, 2, 12, 13 and 17), although the populations, traits and reference assemblies differ enough that this should not be read as replication. Two studies in structured grapevine populations point the same way. Blois et al. (2023) detected 11 QTL for root-related traits in a wild *V. berlandieri* population, accounting for up to 25.1% of the variance, and Morel et al. (2026) found individual QTL explaining only 3.1 to 14.1% of the phenotypic variance for early root-related traits in a *V. berlandieri* × *V. rupestris* population of 449 genotypes. The absence of large-effect loci therefore holds across biparental, single-species and multi-species designs. Morel et al. (2026) add a caution that bears directly on this study: the regions they detected were largely non-overlapping between plants raised from cuttings and grafted plants, so loci mapped from cuttings, as ours are, need not be the loci that matter once a rootstock is grafted and established in the field.

The annotations of genes near the lead SNPs cluster into three processes. The first is auxin-linked cell proliferation and root patterning. The chromosome 8 locus for root weight places the lead SNP inside the grapevine orthologue of *PASTICCINO1*, an FKBP-type peptidyl-prolyl isomerase required for auxin-dependent control of cell division and differentiation and shown to promote root growth when overexpressed (Liu et al. 2024; Pérez-Pérez et al. 2004). The strongest root weight locus, on chromosome 10, carries *SCARECROW-LIKE 21*, a GRAS-family transcription factor of the SHORT-ROOT/SCARECROW module that specifies radial patterning in the root, together with a WRKY transcription factor. An AP2/ERF-*RAV1* transcription factor sits at the chromosome 14 locus; AP2/ERF factors integrate age and wound signals during root regeneration (Ye et al. 2020). At the post-transplant locus on chromosome 11 the nearest genes include a *rac*-like ROP GTPase, a family that controls polar cell expansion and root hair initiation.

The second process is cell wall and lignin remodeling. The callus-stage lead SNP on chromosome 2 lies 30 bp from a mitogen-activated protein kinase and 0.9 kb from an O-acetyltransferase of the family that acetylates cell wall polysaccharides and thereby alters wall flexibility and expansion, a modification relevant to the emergence of root primordia (Zhong et al. 2020). Expansin A1 and a galacturonosyltransferase fall within the core intervals of two root weight loci. The most compositionally distinctive locus of the study belongs to the transplant-response index: on chromosome 11 at 2.04 Mb the lead SNP lies inside one of four caffeoyl-CoA O-methyltransferase genes arrayed within 18 kb. CCoAOMT catalyzes a committed step of monolignol biosynthesis, and reduced lignification has been associated with improved adventitious root emergence (Rogers & Campbell 2004; Chang et al. 2023). The transplant-response index was defined before any association analysis as the change in rooting score across transplanting, and lignification and suberization of the stem base are central to that transition, so the coincidence is worth following up. It remains a hypothesis: the locus needs allelic and expression evidence before the effect can be attributed to *CCoAOMT* rather than to a haplotype that happens to contain those genes.

The third process is solute transport. A tandem array of three OPT oligopeptide transporters sits at the second callus-stage locus, a *NRT1/PTR FAMILY 7.3*-like nitrate transporter within the chromosome 15 root weight locus, a sugar transporter *ERD6*-like 6 at the chromosome 4 locus, and a Major Facilitator Superfamily transporter carries the post-transplant lead SNP. Nitrate transporters of the NPF/NRT1 family link nutrient sensing to auxin transport and lateral root development (Ho et al. 2009; Krouk et al. 2010; Wang et al. 2009), and sugar carriers have been associated with rooting through carbohydrate delivery to the rooting zone (Druege et al. 2019; Husen & Pal 2007; Ruedell et al. 2013). All of these assignments are positional. No expression or functional evidence is presented here, the panel spans 18 species so a lead SNP may tag a haplotype block differentiated between species rather than a causal variant, and the ±25 kb core still contains five genes on average.

Prediction was moderately accurate and, importantly, insensitive to marker number. Accuracy for callus-stage rooting within 2025 rose only from 0.620 to 0.639 when marker number increased tenfold from 5,000 to 50,000, and the same pattern held for every trait and scenario. Accuracy as a function of training-set size was likewise close to flat between 50 and 200 accessions. Neither denser genotyping nor a modestly larger training set is therefore the efficient route to better prediction in this system, which matters because both are the usual first response to disappointing accuracy. Adding spectral information did help: the combined genomic-plus-spectral model was the best model for post-transplant rooting (0.701 against 0.668), root weight (0.605 against 0.580) and the transplant-response index (0.634 against 0.592). A taxonomy-only model, which uses nothing but species identity, reached r = 0.55 for callus-stage rooting. For pre-a taxonomy-only model is useful and costs nothing; its limit is that it cannot rank accessions within a species, and genomic or spectral data are what make within-species selection possible.

These accuracies are best read against what prediction has achieved elsewhere. Grapevine improvement has relied largely on marker-assisted selection for traits under simple genetic control, such as Pierce’s disease resistance at PdR1 (Riaz et al. 2009) and dagger nematode resistance at XiR1 (Hwang et al. 2010), an approach that does not transfer to a quantitative trait like rooting. Genomic prediction was developed for that case (de Los Campos et al. 2013) and has since been evaluated across a wide range of crops and trait types, including wheat (Heffner et al. 2011; Norman et al. 2018), maize (Rio et al. 2019), sorghum (Sapkota et al. 2020), pea (Burstin et al. 2015), cassava (Phumichai et al. 2022), alfalfa (Medina et al. 2021), eucalyptus (Muller et al. 2017), apple (Roth et al. 2020; Minamikawa et al. 2024), pear (Sun et al. 2024) and blueberry (de Bem Oliveira et al. 2020; Cromie et al. 2025). In grapevine it has been applied mainly to scion traits, in table grapes (Viana et al. 2016) and across *V. vinifera* populations (Brault et al. 2022; Brault et al. 2024). The accuracies we obtain for rooting, 0.58 to 0.67 within a season and 0.49 to 0.65 for accessions the model had not seen, are within the range that has been judged useful in those systems, and they are obtained in a panel spanning 18 species rather than within a single breeding population. To our knowledge this is the first application of genomic and phenomic prediction to rootstock traits in wild *Vitis* germplasm.

Climate of origin also carries information. The association between low dry-season precipitation at the collection site and better rooting persisted after species means were removed, and was strongest for root weight. We do not propose a mechanism, and the association is observational, but collection-site coordinates are already recorded for most accessions in this and comparable collections, so the information is free to use as a prior for germplasm that has not yet been evaluated.

Five limitations bear on how these results should be read. First, only one observation per accession per season is available, so genetic and error variance within a season are not separately identifiable and every heritability-like quantity reported here is a repeatability on an entry-mean basis. Second, the prediction accuracies come from random cross-validation in a panel with strong population structure; this is the design the genomic-selection literature reports and is therefore comparable with other studies, but it is optimistic, and the numbers should be read as prediction among relatives of evaluated material rather than as prediction into a new species. Third, the 2026 season behaved differently from the other two on every measure we examined (repeatability, between-season correlation, kappa agreement, and predictability), and that season was scored once per bundle rather than per plant; results involving it are reported with that caveat throughout, and the contrast is the strongest evidence in the study that phenotyping protocol, not marker density or panel size, is the binding constraint. Fourth, the association analysis spans 18 species, and the kinship matrix and principal components control the resulting inflation well (λ = 0.90-1.08) but cannot distinguish a causal allele from a species-diagnostic one. Fifth, every accession was aligned and called against a single haplotype of one genome. Reference bias grows with divergence from *V. girdiana*, so variant discovery is expected to be least complete in the species furthest from it, and sequence absent from the reference haplotype cannot be tested at all; loci may therefore have been missed, but those reported here are not artifacts of the reference choice. A super-pangenome built from this same panel is available (Cochetel et al. 2023), and repeating the analysis against it would settle the question directly. Pangenome-scale association analysis remains computationally demanding at this panel size, but it is a natural next step rather than an obstacle to the present analysis.

Taken together, these results establish that useful rooting variation exists well outside the *riparia*-*rupestris*- *berlandieri* group, that it is heritable enough to select on, that it is predictable at moderate accuracy from a few thousand markers, and that the loci underlying it point to auxin signaling, cell wall and lignin remodeling, and solute transport. Accessions identified here as strong rooters should now be characterized for the traits that rootstocks are actually chosen for, such as nematode resistance, drought and salinity tolerance, and vigor control, so that rooting ability can be combined with them rather than traded against them.

## Data Availability Statement

All data collected in this study are provided in Supplemental Tables S1 to S6 accompanying this manuscript. Raw sequencing reads are available from the NCBI Sequence Read Archive under BioProjects PRJNA731597, PRJNA842753, and PRJNA984685.

## Author Contributions

LDG and SS: conceptualization; LDG: supervision; SS and LDG: writing; LDG: funding acquisition; SS, YL, JM, VN, ETL, and AG: experimentation; SS, NC, DC, and LDG: data analysis and visualization

## Acknowledgments

The authors thank Mikayla Bailey, David Sweet, Gianna Smooth, and Guillermo Garcia-Zamora, for their support with greenhouse experiments and vineyard maintenance.

## Supplemental Material

Supplemental Figure S1. Season-to-season agreement for the two ordinal rooting scores. Supplemental Table S1. Measurement basis by season and trait.

Supplemental Table S2. All 315 genes within ±100 kb of a lead SNP, with coordinates, strand, distance and orientation relative to the lead SNP, a flag marking the ±25 kb linkage-disequilibrium core, and the functional annotation.

Supplemental Table S3. All 85 trait × climate associations with within-species slopes, across-panel correlations, and BH-adjusted q-values.

Supplemental Table S4. Prediction accuracy by spectral preprocessing pipeline. Supplemental Table S5. Prediction accuracy as a function of training-set size.

Supplemental Table S6. Full GWAS summary statistics per trait (λ, SNPs tested, significant SNPs, independent loci, threshold, accessions analyzed, and PVE ± SE from the null model).

## Abbreviations

ARF: adventitious root formation
BH: Benjamini–Hochberg
BLUP: best linear unbiased prediction
GBLUP: genomic best linear unbiased prediction
GWAS: genome-wide association study
LD: linkage disequilibrium
PC: principal component
PVE: proportion of phenotypic variance explained
QTL: quantitative trait locus
REML: restricted maximum likelihood
SNP: single-nucleotide polymorphism
SNV: standard normal variate.

---

**Supplemental Figure S1.**
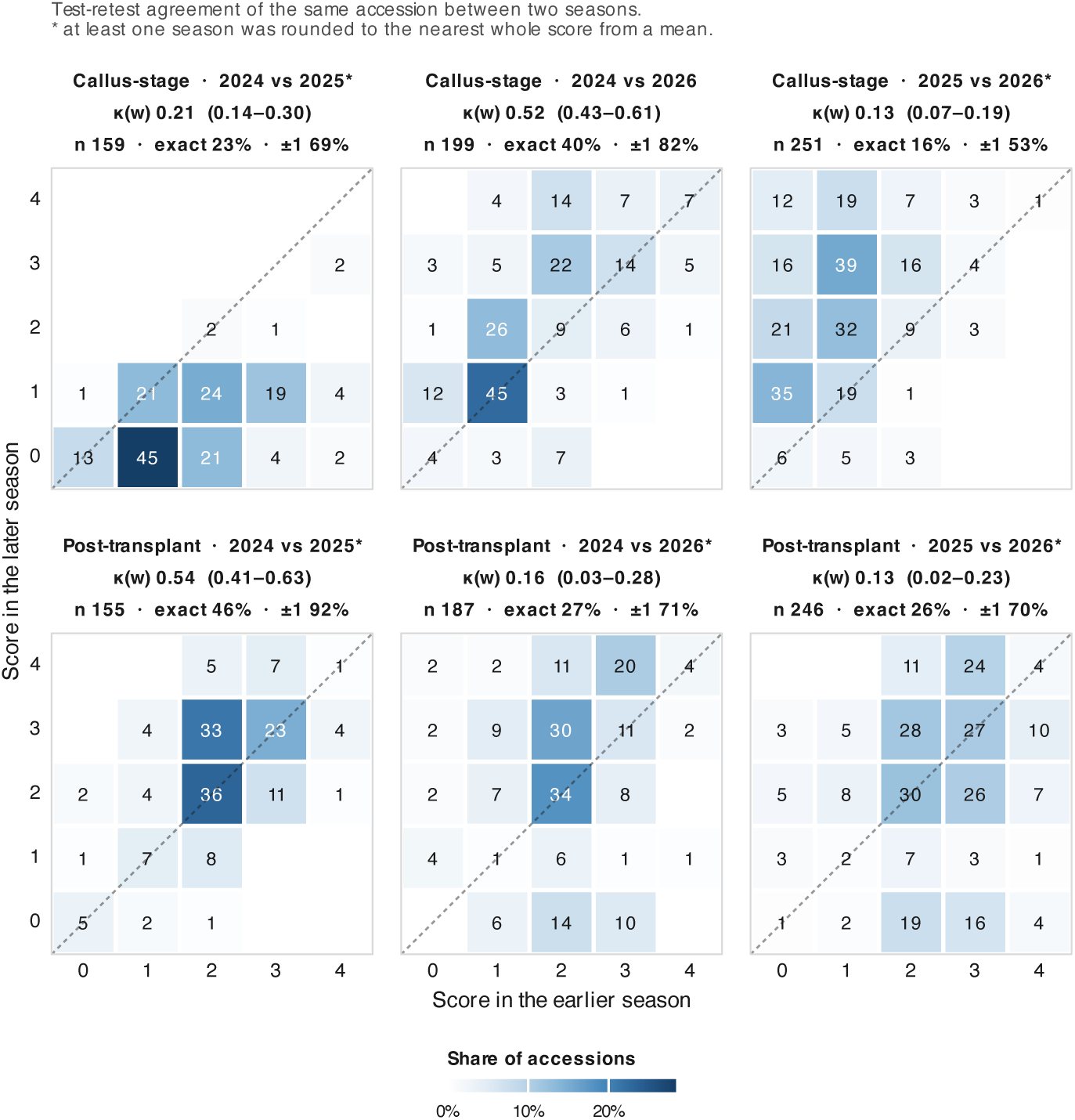
Season-to-season agreement for the two ordinal rooting scores. Test-retest matrices between seasons for the same accession, with quadratic-weighted Cohen’s kappa and 1,000 replicate bootstrap intervals. Comparisons involving a season whose values are means rather than native integers were rounded to the nearest whole score and are marked with an asterisk; the Spearman correlation on the unrounded values is given for comparison.

