## Supplemental Figure 1 for "Harnessing *Vitis* diversity to dissect and predict adventitious rooting traits in grapevine"

Sharma et al.

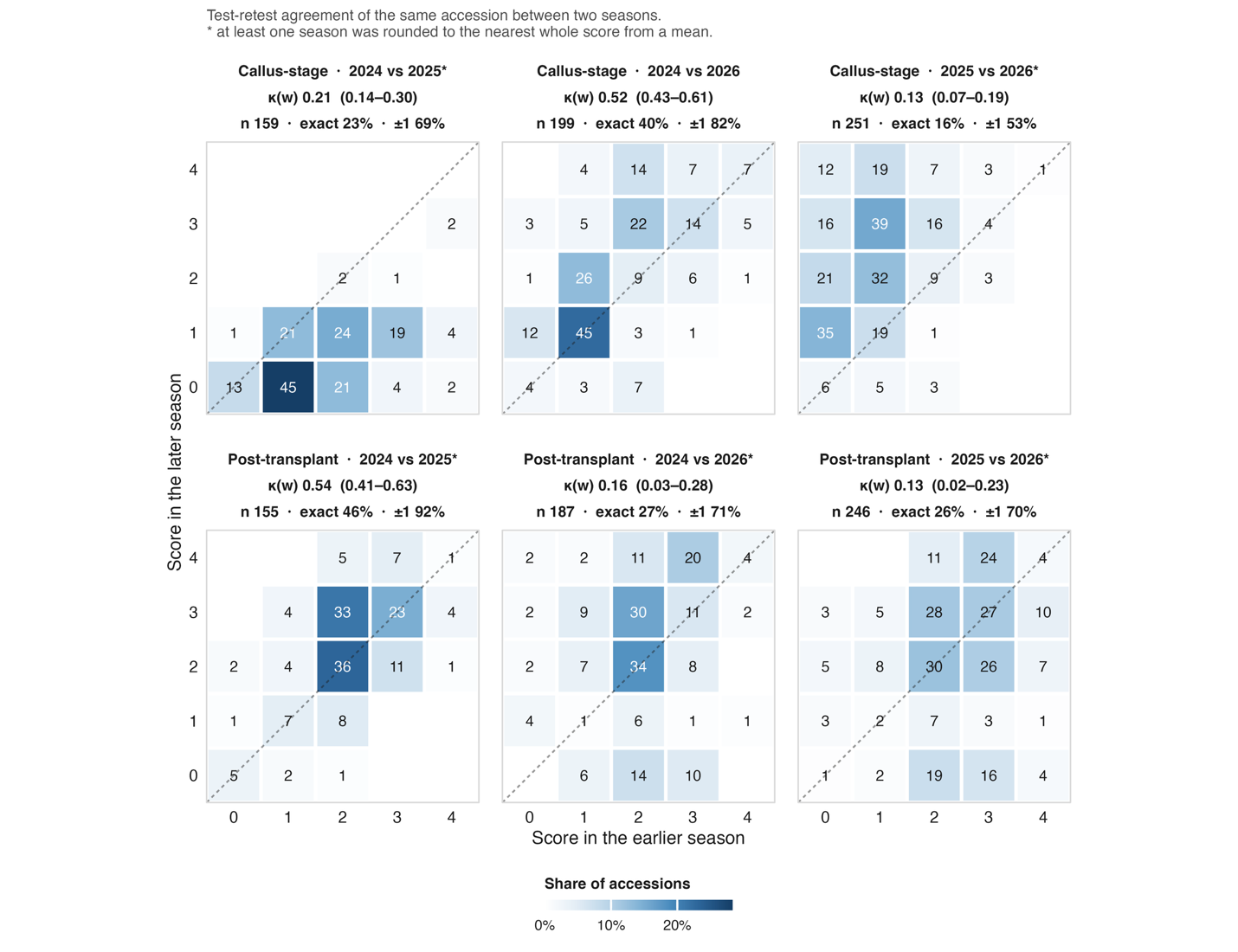

Supplemental Figure S1. Season-to-season agreement for the two ordinal rooting scores. Test-retest matrices between seasons for the same accession, with quadratic-weighted Cohen's kappa and 1,000 replicate bootstrap intervals. Comparisons involving a season whose values are means rather than native integers were rounded to the nearest whole score and are marked with an asterisk; the Spearman correlation on the unrounded values is given for comparison.

Supplemental Tables S1 to S6 are provided as a separate Microsoft Excel workbook, one table per sheet.
